# Resistance gene discovery and cloning by sequence capture and association genetics

**DOI:** 10.1101/248146

**Authors:** Sanu Arora, Burkhard Steuernagel, Sutha Chandramohan, Yunming Long, Oadi Matny, Ryan Johnson, Jacob Enk, Sambasivam Periyannan, M. Asyraf Md Hatta, Naveenkumar Athiyannan, Jitender Cheema, Guotai Yu, Ngonidzashe Kangara, Sreya Ghosh, Les J. Szabo, Jesse Poland, Harbans Bariana, Jonathan D. G. Jones, Alison R. Bentley, Mick Ayliffe, Eric Olson, Steven S. Xu, Brian J. Steffenson, Evans Lagudah, Brande B. H. Wulff

**Author notes:** These authors contributed equally to this work (Sanu Arora, Burkhard Steuernagel).

## Abstract

Genetic resistance is the most economic and environmentally sustainable approach for crop disease protection. Disease resistance (R) genes from wild relatives are a valuable resource for breeding resistant crops. However, introgression of R genes into crops is a lengthy process often associated with co-integration of deleterious linked genes^1, 2^ and pathogens can rapidly evolve to overcome R genes when deployed singly^3^. Introducing multiple cloned R genes into crops as a stack would avoid linkage drag and delay emergence of resistance-breaking pathogen races^4^. However, current R gene cloning methods require segregating or mutant progenies^5–10^, which are difficult to generate for many wild relatives due to poor agronomic traits. We exploited natural pan-genome variation in a wild diploid wheat by combining association genetics with R gene enrichment sequencing (AgRenSeq) to clone four stem rust resistance genes in <6 months. RenSeq combined with diversity panels is therefore a major advance in isolating R genes for engineering broad-spectrum resistance in crops.

---

Plant disease resistance is often mediated by dominant or semi-dominant R genes which encode receptors that perceive pathogen infection either by direct recognition of pathogen effector molecules, or indirectly by recognition of effector modified host targets^11^. Most known R genes encode intracellular nucleotide binding/leucine-rich repeat (NLR) immune receptor proteins^11^. NLR genes are amongst the most evolutionarily dynamic gene families in plants with hundreds of these genes usually present in plant genomes^12^. Crop plants and their wild progenitors typically possess extensive copy number and sequence variation at these NLR loci^12, 13^. Domestication and intensive breeding have reduced R gene diversity in crops rendering them more vulnerable to disease epidemics^14^. New R genes derived from closely related species have been introduced into crops to improve disease resistance by lengthy and laborious introgression breeding or, more recently, as transgenes^10, 15–17^.

R gene cloning, as needed to introduce exotic R genes, is complicated by extensive pan-genome variation at NLR loci making it difficult to establish orthogonal relationships between a reference genome sequence and other accessions^13^. Structural variation at NLR loci of different accessions can also reduce recombination at a locus^18^ further complicating positional cloning approaches. The requirement for recombination in gene cloning can be overcome by mutational genomics. The generation of multiple independent mutants by mutagenesis followed by genome complexity reduction and sequencing (e.g. NLR exome capture (MutRenSeq^5^) or chromosome flow sorting and sequencing (MutChromSeq^6, 19^)), enables unambiguous identification of mutant alleles and the functional NLR gene (Table 1).

**Table 1.**
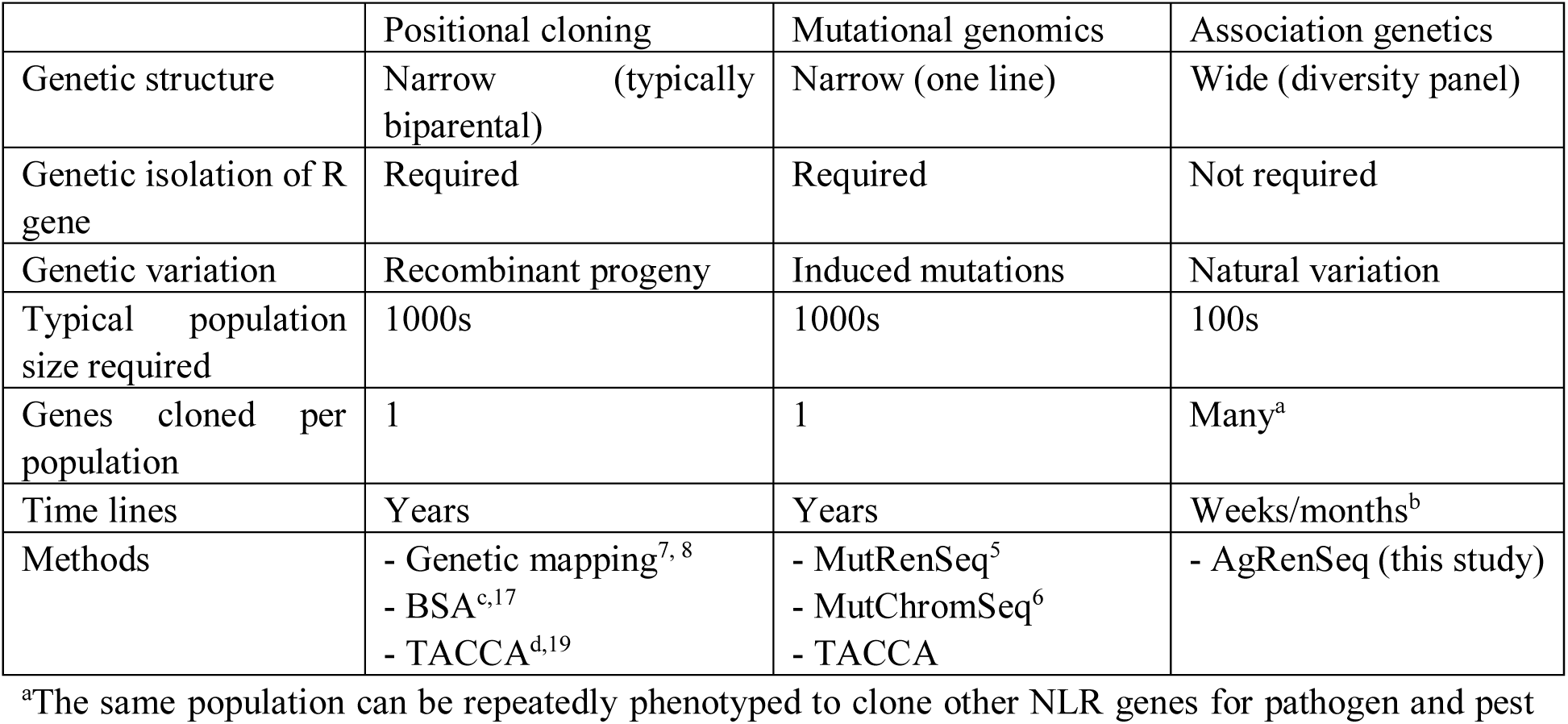
Comparison of R gene cloning strategies.

|  | Positional cloning | Mutational genomics | Association genetics |
| --- | --- | --- | --- |
| Genetic structure | Narrow (typically biparental) | Narrow (one line) | Wide (diversity panel) |
| Genetic isolation of R gene | Required | Required | Not required |
| Genetic variation | Recombinant progeny | Induced mutations | Natural variation |
| Typical population size required | 1000s | 1000s | 100s |
| Genes cloned per population | 1 | 1 | Many<sup>a</sup> |
| Time lines | Years | Years | Weeks/months<sup>b</sup> |
| Methods | - Genetic mapping<sup>7, 8</sup><br>- BSA<sup>c,17</sup><br>- TACCA<sup>d,19</sup> | - MutRenSeq<sup>5</sup><br>- MutChromSeq<sup>6</sup><br>- TACCA | - AgRenSeq (this study) |

<sup>a</sup>The same population can be repeatedly phenotyped to clone other NLR genes for pathogen and pest resistance.
<sup>b</sup>Only phenotyping is required once the diversity panel has been configured.
<sup>c</sup>Bulked segregant analysis
<sup>d</sup>Targeted chromosome-based cloning via long-range assembly.

Positional cloning and mutational genomics share certain limitations. Notably, they require the R gene to exist as a single gene in an otherwise susceptible genetic background to the pathogen isolate of interest which can take numerous generations to achieve. They also require that the plant target species is amenable for the generation and screening of large recombinant or mutant populations numbering 1000s of individuals (Table 1). However, many wild relatives of crop plants contain multiple functional resistance genes and are experimentally intractable due to pre-domestication traits such as long generation times, seed shattering, and seed dormancy, as well as having seed and plant morphologies not amenable to existing mechanized technologies.

In contrast, genome-wide association studies (GWAS) do not require monogenic resistant lines or the development of mapping or mutagenesis populations. GWAS enables trait correlation in a genetically diverse population structure by exploiting pre-existing recombination events accumulated in natural populations. Members within a population are screened with a genome-wide set of markers and linkage disequilibrium sought between markers and traits of interest.

Markers linked to traits of interest are then readily assigned to a genomic region in a reference genome^20^. However, the reliance on a reference genome complicates the identification of sequences that have significantly diverged from the reference, as is often the case for R gene loci. Recently, this limitation was overcome by performing trait associations on sub-sequences (k-mers) different between case and control samples to identify variants associated with human disease^21^ or bacterial antibiotic resistance^22^. We reasoned that k-mer–based association genetics combined with resistance gene enrichment sequencing (AgRenSeq) would enable the discovery and cloning of R genes from a plant diversity panel (Fig. 1).

**Figure 1.**
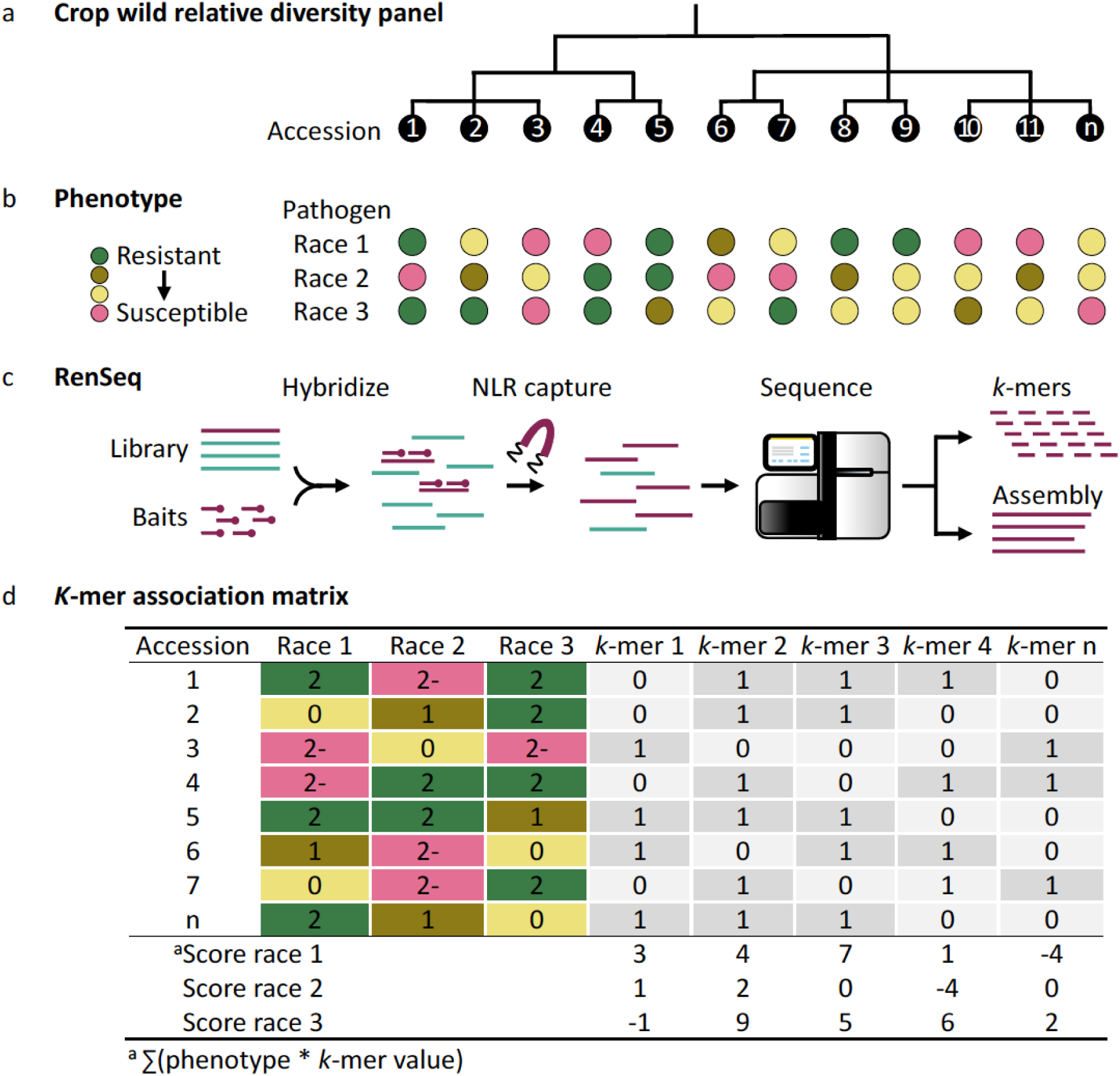
Combining association genetics and resistance gene enrichment sequencing (AgRenSeq) for rapid R gene discovery and cloning. (a) A genetically diverse panel of accessions is (b) phenotyped with different pathogen races, and (c) subjected to RenSeq followed by assembly of the NLR repertoire and extraction of NLR k-mers for each accession. (d) Each k-mer in each accession was given a phenotype score between +2 and -2 depending on the level of resistance or susceptibility. The sum of scores for each k-mer is calculated for each pathogen race. k-mers are then plotted in an association matrix according to their sequence identity to NLRs from a given accession (x-axis) and their phenotype score (y-axis) (see Fig. 2).

Here, we describe AgRenSeq analysis of a panel of Aegilops tauschii accessions that were phenotyped with races of the wheat stem rust pathogen Puccinia graminis f. sp. tritici (PGT). Ae. tauschii is the wild progenitor species of the Triticum aestivum (bread wheat) D genome and a valuable source of stem rust resistance (Sr) genes that have been introgressed into bread wheat. To test AgRenSeq, we chose Ae. tauschii ssp. strangulata because of the availability of genetically diverse accessions^23^ and two cloned Sr genes from this species (Sr33 and Sr45) to serve as positive controls^5, 7^. We obtained 174 Ae. tauschii ssp. strangulata accessions that span its habitat range (Supplementary Table 1) and also included 21 Ae. tauschii ssp. tauschii accessions as an outgroup.

We designed and tested a sequence capture bait library optimised for Ae. tauschii NLRs plus genomic regions encoding 317 single nucleotide polymorphism (SNP) markers distributed across all seven chromosomes (Supplementary Fig. 1; Supplementary Tables 2-3). RenSeq analysis was undertaken on the diversity panel using this capture library and Illumina short-read sequencing, with an average of 1.67 Gb of 250 bp paired-end sequence obtained per accession (Supplementary Table 4). De novo sequence assemblies were analysed with NLR-parser, a program that identifies NLRs based on number and order of short conserved NLR sequence motifs^24^. We obtained between 1,312 and 2,170 (average 1,437) NLR contigs per accession, of which 249 to 336 (average 299) encoded full length NLRs and 1,024 to 1,921 (average 1,137) encoded partial NLRs (Supplementary Table 4). Captured SNP marker sequences were compared among accessions and a set of 151 genetically distinct Ae. tauschii ssp. strangulata accessions were selected (Fig. 2a; Supplementary Fig. 2-3; Supplementary Table 5). These accessions were phenotyped for their reaction to seven PGT races with different virulence profiles and geographic origins (Supplementary Table 6; Supplementary Fig. 4. From 10 to 67% of the accessions showed resistance depending on the PGT race used (Fig. 2b; Supplementary Fig. 5; Supplementary Table 7).

**Figure 2.**
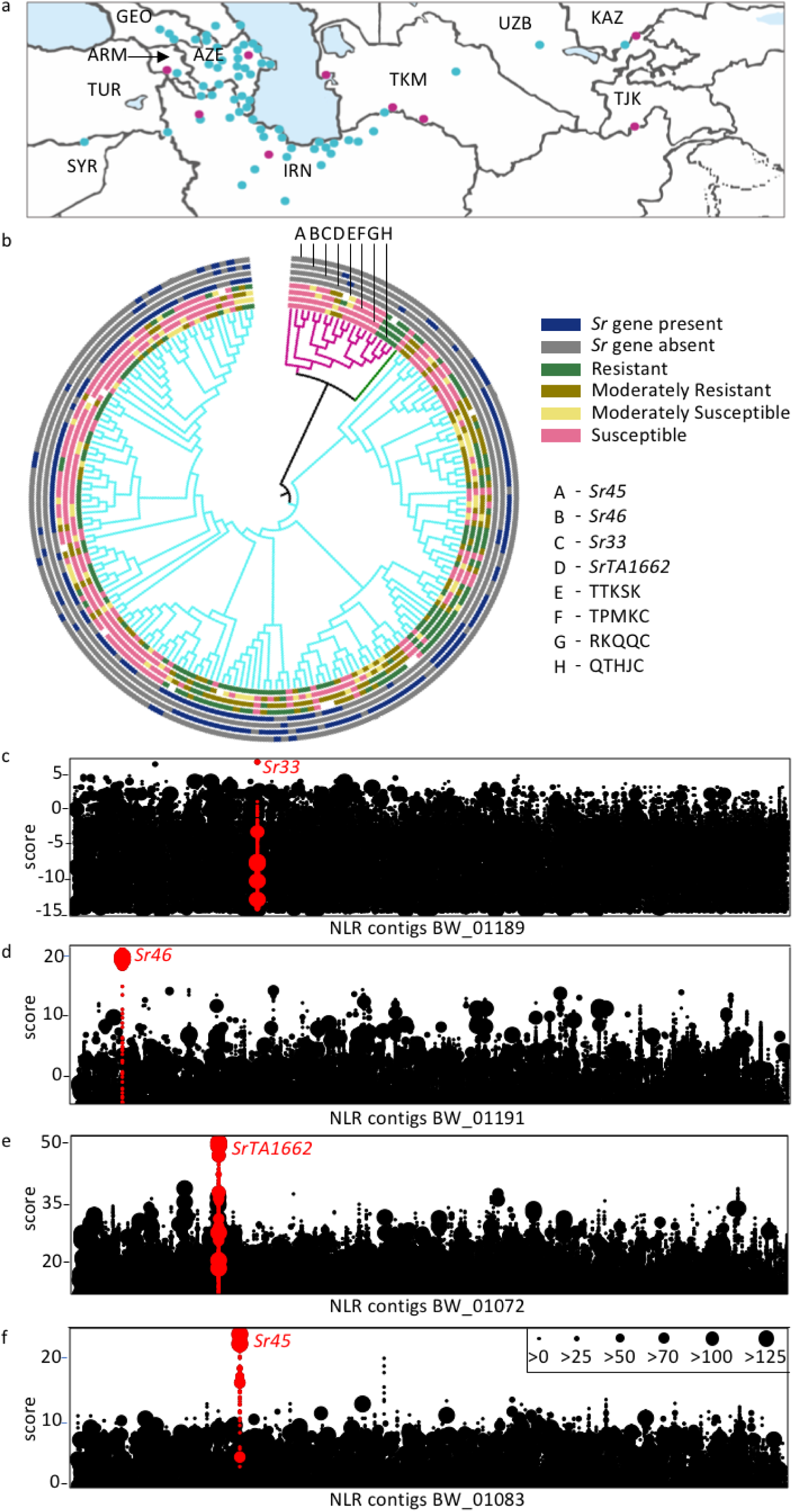
Genetic architecture of stem rust resistance in Ae. tauschii. (a) Geographic distribution of Ae. tauschii ssp. strangulata (cyan) and ssp. tauschii (magenta) used in this study. Two accessions from China and two from Pakistan, which fall outside of the map, are not shown. ARM, Armenia; AZE, Azerbaijan; GEO, Georgia; IRN, Iran; KAZ, Kazakhstan; SYR, Syria; TJK, Tajikistan; TUR, Turkey; TRK, Turkmenistan; UZB, Uzbekistan (b) Phylogenetic tree displaying Ae. tauschii ssp. strangulata (151 accessions, cyan) and ssp. tauschii (21 accessions, magenta) with an intermediate accession (dark green). Stem rust phenotypes and Sr genotypes are displayed by concentric circles around the tree. (c-f) Identification of Sr33, Sr46, SrTA1662 and Sr45 by AgRenSeq using PGT races RKQQC, TPMKC, QTHJC and TTKSK, respectively. Each dot-column on the x-axis represents an NLR contig from the RenSeq assembly of a single accession containing the respective Sr gene. Each dot on the y-axis represents one or more RenSeq k-mers associated with resistance across the diversity panel to the respective PGT race. Dot size is proportional to the number of k-mers associated with resistance.

To identify stem rust resistance (Sr) genes in the phenotyped panel, we performed a k-mer– based AgRenSeq analysis. The k-mer sequences were given a numerical value based on the level of resistance or susceptibility observed for the accession in which they occurred, and these values were then summed up for each k-mer. In previous k-mer based GWAS, associated k-mers were used to build a local assembly^21^ or reconstruct diverged haplotypes through a direct base-by-base k-mer frequency-guided extension process^22^. However, we reasoned that local assembly approaches using only those k-mers that are strongly linked to the trait would not generate complete NLR contigs especially across the more conserved NB-ARC (nucleotide-binding adaptor shared by APAF-1 R proteins, and CED-4) domain. We therefore projected k-mers directly onto NLR assemblies generated from read data of resistant accessions to obtain longer contigs including full length NLRs.

A proof-of-concept experiment was undertaken to identify the Sr33 gene amongst these accessions using North American PGT race RKQQC, which is avirulent to Sr33, but virulent to most other known Sr genes^25^. We mapped k-mers associated with RKQQC resistance onto the RenSeq assembly of accession BW_01189 which is the original source of Sr33^7^. A single discrete association peak was identified in a 5.6 kb RenSeq contig that had 100% identity to Sr33 (Fig. 2c). The same method was then used to map k-mers associated with resistance to all six remaining PGT races in the 151 accessions. Two non-redundant, high-confidence candidate Sr genes other than Sr33 were identified (Fig. 2d-e, Supplementary Table 7) and these candidate Sr genes were aligned to the reference genome assembly of wheat cv. Chinese Spring RefSeq1.0 (www.wheatgenome.org). The genomic locations of these candidate Sr genes coincided with the genetic positions of two Sr genes introgressed into wheat from Ae. tauschii ssp. strangulata, namely Sr46^26^ and SrTA1662 ^27^. The SrTA1662 candidate gene was mapped in a recombinant inbred line population to a 3.8 cM interval shown to encode SrTA166-resistance, thus supporting our AgRenSeq data (Supplementary Fig. 6; Supplementary Table 8). The SrTA1662 candidate encodes a coiled-coil NLR protein with 85% amino acid identity to Sr33.

Independently of this AgRenSeq approach, we also identified an Sr46 candidate gene by conventional fine mapping in segregating diploid progenitor and wheat populations coupled with the sequencing of candidate genes in this region in three ethyl methanesulphonate-derived mutants that had lost Sr46 resistance. In two mutants, the same candidate gene contained non-synonymous substitutions, while the third mutant had a deletion of the chromosomal segment encoding this gene (Supplementary Fig. 7a-f). Comparison of the Sr46 candidate gene identified by AgRenSeq and map-based/mutagenesis cloning showed that they were 100% identical. This gene, hereafter referred to as Sr46, encodes a coiled coil NLR protein that conferred stem rust resistance when expressed as a transgene in the susceptible wheat cv. Fielder (Supplementary Fig. 7g, Supplementary Table 9).

While identification of Sr33, Sr46 and SrTA1662 was successful, we did not identify Sr45. We hypothesized this was due to the prevalence of Sr46 in the panel thereby masking Sr45 resistance as none of the seven PGT races utilized were virulent to Sr46 and avirulent to Sr45. To overcome this constraint, we removed all 64 accessions containing Sr46 from the panel. The k-mers associated with resistance to PGT race TTKSK in the remaining 87 accessions were mapped onto the TTKSK-resistant Ae. tauschii accession BW_01083, the original donor of Sr45^5^. Previously we identified a candidate Sr45 gene by MutRenseq from a wheat-Ae. tauschii BW_01083 introgression line^5^. Using AgRenSeq, a single discrete association peak was identified that corresponded to this previously identified Sr45 candidate gene (Fig. 2f). This gene conferred resistance when transformed into cv Fielder confirming both our previous MutRenSeq identification^5^ and current AgRenSeq data (Supplementary Fig. 8, Supplementary Table 9). From AgRenSeq analysis and sequence alignments among these four cloned Sr genes and the diversity panel, Sr46 and SrTA1662 were the most prevalent being found in 42% of the accessions, while Sr33 and Sr45 were found in 5% and 7% of accessions, respectively (Fig. 2b).

Our results demonstrate that AgRenSeq is a robust protocol for rapidly discovering and cloning functional R genes from a diversity panel. This approach successfully identified four Sr genes that have been introgressed into wheat from Ae. tauschii ssp. strangulata over the last 40 years^5, 7, 26, 27^. In addition to stem rust resistance, Ae. tauschii is a rich source of genetic variation for resistance to other diseases and pests relevant to bread wheat cultivation including leaf rust, stripe rust, wheat blast, powdery mildew, Hessian fly, and others (Supplementary Table 10). Our RenSeq-configured diversity panel is essentially an R gene genotyped resource that can now be screened against other pathogens/pests to clone additional functional R genes.

In contrast to most published association studies, the AgRenSeq procedure is reference genome-independent and directly identifies the NLR that confers resistance rather than identifying a genomic region encoding multiple paralogs for subsequent candidate gene confirmation. AgRenSeq is also not restricted to the limited genetic variation and recombination present in bi-parental populations, but rather interrogates pan-genome sequence variation in diverse germplasm to isolate uncharacterised R genes. Thus, importantly, no additional crossing or mutagenesis is required for R gene cloning using this approach. This makes it possible to clone R genes from accessions of a wild species lacking any advanced agronomic traits and requires only the phenotyping of an enrichment-sequenced diversity panel.

Germplasm collections are available for many crops and their wild relatives including soybean, pea, cotton, maize, potato, wheat, barley, rice, banana and cocoa (Supplementary Table 11). The ability to rapidly clone agriculturally valuable R genes by AgRenSeq further increases the value of these germplasm collections. AgRenSeq may also allow breeders to discover NLR genes in their breeding germplasm and develop gene-specific markers for marker-assisted selection. Thus, this method has enormous utility for genetic improvement of most crops and will also provide new fundamental insights into the structure and evolution of species-wide functional R gene architectures.

Accession codes. Sequence reads were deposited in the European Nucleotide Archive (ENA) under project number X. The Sr46 and SrTA1662 loci were deposited at NCBI under accession numbers X and Y. The programs and scripts used in this analysis have been published on Github (https://github.com/steurnb/XXX/ and https://github.com/aroras/XXX). The bait library sequences are available from https://github.com/steurnb/XXX/. Ae. tauschii accessions with the GRU accession numbers in Supplementary Table 1 are available from the Germplasm Resources Unit, John Innes Centre, Norwich, UK (https://www.jic.ac.uk/germplasm/).

## ACKNOWLEDGMENTS

We are grateful to the germplasm banks at Kansas State University, International Center for Agricultural Research in the Dry Areas, USDA-ARS National Small Grains Collection, and the N. I. Vavilov Research Institute of Plant Industry, and to our colleagues Jan Dvorak and Colin Hiebert for providing seed of Ae. tauschii. We thank our colleagues Yajuan Yue and JIC Horticultural Services for plant husbandry, James Brown for helpful discussions, Mike Ambrose and Alicia Meldrum for help with MTAs, Kamil Witek for technical assistance with RenSeq, the International Wheat Sequencing Consortium for pre-publication access to REFSEQ v1.0, and Bob McIntosh for critical reading of the manuscript. This research was supported by the NBI Computing Infrastructure for Science (CiS) group and financed by the Two Blades Foundation, USA to BJS and BBHW; the Biotechnology and Biological Sciences Research Council (BBSRC) to BBHW and the BBSRC Designing Future Wheat Cross-Institute Strategic Programme to ARB and BBHW, the Norwich Research Park Translational Fund to BBHW; the Lieberman-Okinow Endowment at the University of Minnesota to BJS; the Australian Grains Research and Development Corporation to EL and HB; and in-kind support by Arbor Biosciences.

## AUTHOR CONTRIBUTIONS

SA, ARB, JP, SG, EL and BBHW configured diversity panel. SA and JC extracted DNA and performed phylogenetic analysis. BS, SA and JE designed and tested bait library. OM, RJ, SA, NK, LJS and BJS phenotyped diversity panel. JE prepared enriched libraries. SA, GY, BS and BBHW performed AgRenSeq pilot studies. BS designed, implemented and visualised AgRenSeq data correlation matrix. SC, YL, SP, NA, HB, MA, SX, and EL map-base cloned Sr46. SP, MAMdH and MA made Sr45 transgenics. EO mapped SrTA1662. BBHW conceived study and drafted manuscript with SA, BS, JDGJ, ARB, MA, EO, SX, BJS and EL. All authors read and approved the final manuscript.

